# Lightweight Adaptive and Recoverable Implants (LARI) for Chronic Single and Dual Neuropixels 2.0 Recordings for Naturalistic Behaviors in Feature-Rich Environments

**DOI:** 10.64898/2026.01.13.699139

**Authors:** Maibam R Singh, Christopher Gregg

**Author notes:** Co-Corresponding authors: Maibam R Singh, Christopher Gregg.

## Abstract

Understanding the neural mechanisms underlying naturalistic decision-making requires chronic, high-quality neural recordings from distributed brain regions in freely behaving animals. However, existing silicon probe implants for mice are limited by weight, fragility, lack of independent probe adjustment, and poor suitability for long-term recordings in complex environments. Here we introduce LARI-1.0 (Lightweight Adjustable and Reusable Implant) and LARI-mini, two Neuropixels 2.0 implant systems designed for stable, recoverable, and independently adjustable single- and dual-probe recordings in mice. LARI-1.0 enables independent vertical and lateral positioning of two probes, while LARI-mini provides a lighter single-probe alternative, reducing implant mass by ~20%. Fully assembled weights of ~2.3 g (LARI-1.0) and ~1.5 g (LARI-mini) remain within 10% and 7.5% of adult male mouse body weight, respectively. We validated implant performance in a custom feature-rich naturalistic foraging environment that elicited full-body movements, rapid elevation changes, and ethological foraging behaviors across fed, fasted, and high-fat diet states. Both implants exhibited excellent mechanical stability, minimal motion-related artifacts, and probe drift comparable to previous studies. High-quality single-unit activity with stable waveform features, 5–6× signal-to-noise ratios, and no systematic loss of well-isolated units was maintained for up to 180 days. These results demonstrate that LARI-1.0 and LARI-mini provide lightweight, durable, and customizable implant solutions enabling long-term, high-fidelity neural recordings during naturalistic behavior.

## Introduction

Naturalistic animal behavior consists of a rich repertoire of decisions and actions that emerge from an animal’s interactions with diverse and often unfamiliar environmental variables. In contrast to operant-conditioning paradigms, in which animals are trained over days to weeks to perform a restricted set of task-specific actions, naturalistic behaviors unfold within rich ecological contexts, requiring animals to integrate heterogeneous environmental cues while generating spontaneous, high-dimensional movements, rapid context-dependent decisions, and adaptive learning in real time ^1–4^. In naturalistic settings, animals express native stereotyped and spontaneous actions and make decisions that are not necessarily observed in controlled behavioral tasks ^2,5–7^. The neural mechanisms underlying these behaviors and their modulation by environmental experience and internal state remain poorly understood, partly due to limited tools for studying spontaneous behavior in complex environments. Naturalistic foraging offers a particularly powerful paradigm for probing core principles of brain function ^8,9^, and serves as the focus of our investigations into the genetic and neural bases of decision-making ^10–13^.

Recent advances in high-density *in vivo* electrophysiology now enable high-fidelity recordings from hundreds to thousands of neurons across multiple brain regions in freely moving animals^14–20^. These technologies provide unprecedented opportunities to examine the neural dynamics underlying naturalistic behaviors, rather than the more common head-fixed or simplified lab tasks that restrict movement or limit access to multiple behavioral contexts to test specific hypotheses. However, because high-density silicon probes are lightweight and fragile, achieving stable chronic recordings in freely behaving animals navigating feature-rich environments remains challenging.

Neuropixels 2.0 probes are among the smallest and lightest high-density silicon probes currently available, offering one of the highest channels and electrode counts suitable for chronic recordings^17^. Their ability to target both cortical and deep subcortical regions makes them particularly powerful for studying interactions among distributed neural circuits. However, chronic recordings in complex naturalistic environments require implant fixtures that can protect the delicate probes from lateral impacts, maintain stable positioning for artifact-free single-unit recordings over months, and allow recovery and reuse of the probes^18,20–25^. Without such features, researchers are limited to short-term, non-recoverable implants, which is financially prohibitive for many laboratories.

Although the use of recoverable Neuropixels implants is increasing, only a few multi-probe designs exist for mice and they were tested in simple open-field environments instead of more complex environments with naturalistic features^21–25^. Only a small number of multi-probe implants weigh less than 10% of the average adult mouse’s body weight, and only one existing design currently allows dual Neuropixels 2.0 probe implantation in a single fixture^25^. A recent study reported chronic implantation of six Neuropixels 2.0 probes in one hemisphere of the mouse brain, with a total implant weight of approximately 3.4 grams^22^. The implant design used in the study allowed probes to have independent angles of insertion, however, adding the two head-stages required for the 4 probes in the freely moving fixture would bring the overall weight more than 4 grams. This will push the weight of the fixture close to 20% of an average mouse’s body weight, and will likely affect natural movement. Additionally, performing craniotomies for 4–6 probes and then individual insertion for each probe would greatly increase surgical duration, requiring anesthesia for more than 5–6 hours^22^. An improved solution is needed.

For long-term naturalistic foraging studies, an ideal implant for mice must therefore support simultaneous implantation of 2–3 probes (up to 4 maximum), maintain a total assembly weight—including probes, fixtures, and headstages—below 10% of an adult male mouse’s body weight (approximately 2.5 grams), and permit independent vertical depth and lateral positioning of each probe. The available chronic implant designs for Neuropixels 2.0 doesn’t includes these features. These capabilities are essential when targeting brain regions with different dorsoventral depths, such as the frontal cortex and the hypothalamus or laterally adjacent regions like the retrosplenial and entorhinal cortices. Such flexibility ensures that probes do not inadvertently enter regions outside the target areas, and that all shanks can be positioned optimally within the intended nuclei. The Apollo implant designed by Bimbard et al. ^25^ allows dual Neuropixels 2.0 insertion and has a lightweight footprint (~2.2 gram, including headstage), but it does not support independent depth adjustment or flexible lateral spacing between probes. These limitations complicate targeting of brain regions with different depths or narrow spatial boundaries.

To address these constraints, we developed LARI-1.0 (Lightweight Adjustable and Reusable Implant), a novel implant designed for use with either single or dual Neuropixels 2.0 probes. LARI-1.0 allows independent adjustment of both probe depth and lateral distance, enabling selective targeting of brain regions separated dorsoventrally or laterally while keeping neighboring regions undisturbed. We also designed LARI-mini, a single-probe-only version that reduces implant weight by an additional ~20% relative to LARI-1.0. With one of the lightest implant payloads among recoverable mouse-compatible systems, our designs enable long-duration, high-quality neural recordings from multiple brain regions during naturalistic behavior in a complex, feature-rich environment, while allowing probe reuse with minimal reassembly. Using these implants, we successfully recorded from multiple animals for up to 180 days using both single- and dual-probe configurations targeting distant brain regions, enabling longitudinal investigation of complex naturalistic foraging behavior.

## Results

### Feature-Rich Naturalistic Foraging Environment

To record from foraging mice, we designed a naturalistic assay that includes key features from our published foraging assay, including a home base cage and attached foraging arena with patches of sand with or without seeds^10,12,13^. The design is modified to accommodate the logistics of chronic recordings to allow free unobstructed movement of the data cable and includes complex features that test the stability of LARI-1.0 and LARI-mini-implants. The home cage is connected to a foraging arena through a small opening equipped with an automatic door, accessible via a ladder inside the home cage (**Figure 1A**). Once the door opens, mice can freely transition between the home and foraging environment, mimicking natural scenarios in which an animal leaves its burrow to explore or collect food^26–30^.

**Figure 1.**
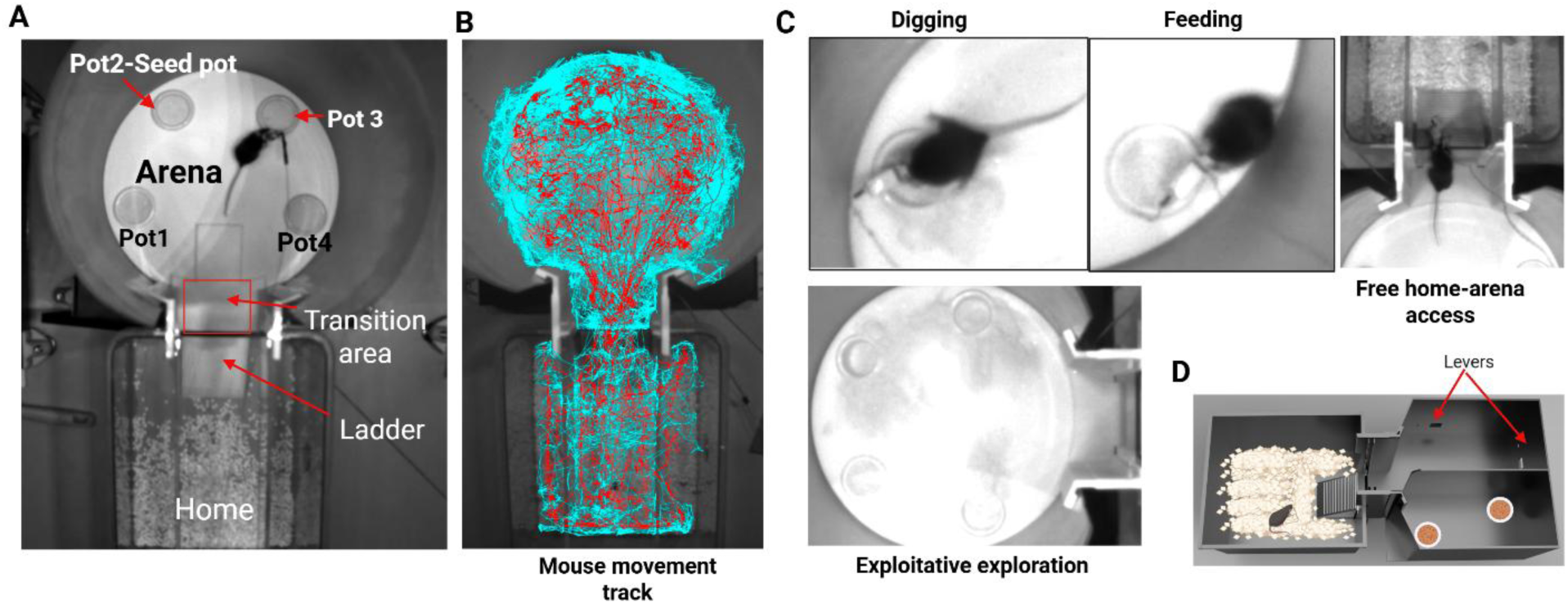
An assay to study the neural computations of naturalistic foraging. **(A)** Naturalistic foraging arena with key features of an ecology setting – foraging arena harboring food patches with sand and seeds is connected to a home cage. The home cage is attached to the arena with a narrow passage guarded by an automatic door, and a ladder to the arena from the home cage. The home cage and the arena have a height difference of 12 cm. Mice have access to both the arena and home throughout the 30-minute “Exploration” and “Foraging” phase sessions. **(B)** Full track of mouse nose and center-point extracted through deep-learning based tracking (Ethovision). **(C)** Naturalistic behaviors such as digging, eating, return to home and explorative behavior are expressed in the naturalistic foraging paradigm. **(D)** We also tested our mouse in another iteration of the foraging arena where mice encountered free access to seed pots, with seeds hidden in one of the pots, but were also given an additional choice to make an effort to acquire reward by pressing one of the levers. This allows one to contrast operant-conditioning style feeding behaviors to more naturalistic foraging.

The foraging arena contains four sand pots (**Figure 1A**). Our foraging assay consists of two phases: (1) a 30-minute Exploration phase, where mice explore a novel environment and find a food patch (Pot 2), and (2) a 30-minute Foraging phase, 4 hours later, where food is relocated and buried (Pot 4). The Exploration phase assesses novelty response, whereas the Foraging phase evaluates memory-driven behavior ^10,12^. Mice perform several different round-trip foraging excursions from the home in this assay ^10^. During the Exploration phase, seeds are placed on top of the sand in Pot2, whereas during the Foraging phase seeds are buried 3.2 cm beneath the sand in one randomly selected pot. The combined area of the home cage (1050 cm²) and the foraging arena (923 cm²) was approximately 2000 cm², with a height difference of 9 cm between the two zones. The elevation difference was intentionally included both to mimic naturalistic vertical transitions and tests whether rapid elevation changes generate data cable induced artifacts.

Each recording day consisted of three phases: 15 minutes of home-only behavior, 30 minutes of Exploration phase, and 30 minutes of Foraging phase, totaling 1 hour and 15 minutes per session. This extended recording duration was chosen because our paradigm did not involve pretraining or habituation, and we aimed to capture a broad repertoire of naturalistic behaviors—including novelty detection, exploration, learning, exploitation, habituation, and excursion-based stereotypical foraging sequences—within a single session. Mice exhibited behaviors such as digging, feeding post digging (**Figure 1C**), and exploratory digging in all sand pots regardless of which contained food (**Figure 1C**, bottom). The feature-rich, multi-planar environment elicited vigorous full-body movements such as digging, rapid darts, and sharp turns, providing a rigorous test of implant stability and spike quality. Alternative versions of the foraging assay were also developed to compare operant versus naturalistic foraging behaviors (**Figure 1D**).

### LARI-1.0 and LARI-mini Neuropixels Recording Systems

To capture robust, longitudinal Neuropixels recordings from foraging mice we developed and tested the LARI-1.0 and LARI-mini implant systems. During pilot recordings, we found that one of the strongest noise sources in the data originated from sudden elevation shifts when mice darted between the home and foraging arena, highlighting the importance of securing the flex cable and headstage to minimize mechanical coupling. Both LARI-1.0 and LARI-mini were specifically designed to suppress these forces by tightly housing the flex cable and head stage within the implant (**Figure 2A-C**).

**Figure 2.**
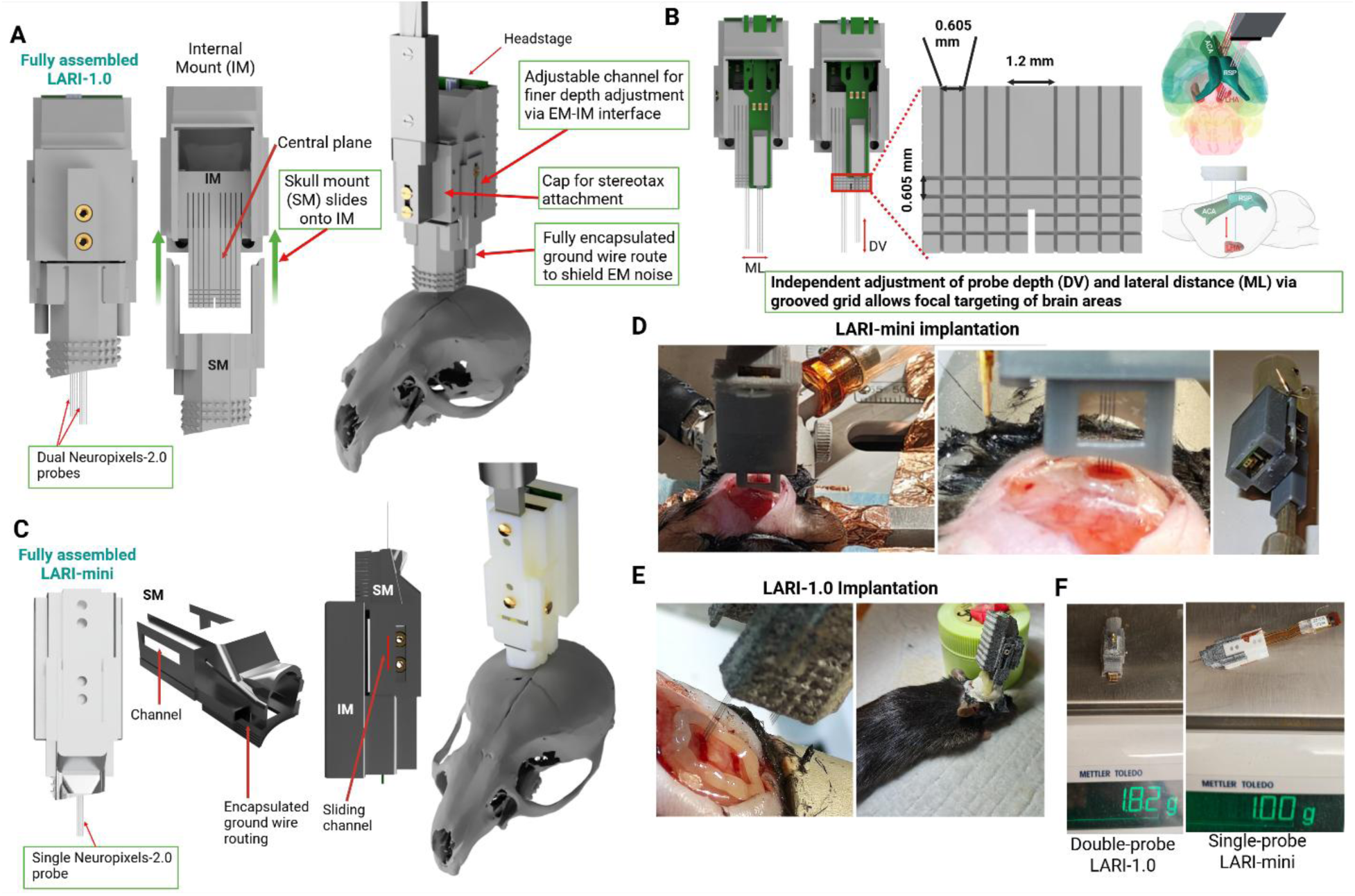
Design and features of the LARI-1.0 and LARI-mini Neuropixels recording systems. **(A)** Components of LARI-1.0 system. The implant fixture consists of an internal mount (IM), a skull mount (SM) that is permanently affixed to the skull, and a Cap that covers the probes and attaches to the stereotax rod. Both faces of the *central plane* in the IM houses one Neuropixels-2.0 probe. A representative illustration of a fully assembled dual probe LARI-1.0 implanted is shown sitting atop a mouse skull *(far right)*. There’s dedicated protective sleeve in the SM and IM through which the ground cable is routed. This reduces ground wire exposure and ensures shielding of external module (EM) noise. |**(B)** Central plane in LARI-1.0 IM with marked and equally spaced (0.605 mm) vertical and horizontal grooves for independent adjustment of relative depth and lateral distance between the two Neuropixels-2.0 probes. Four holes of different sizes are provided on the IM, two for brass inserts and the other two for use with M1.2 screws without the need of inserts, to attach the SM. This allows further fine adjustment of the effective shank depth via custom positioning of SM. **(C)** LARI-mini fixture for single Neuropixels-2.0 probe implantation only consists of the SM and IM. The IM consists of an integrated sleeve to attach to a custom stereotax rod. Similar to dual-probe LARI, effective depth of the probe can be adjusted via both the IM and EM. **(D** and **E)** Shows surgical implantation of LARI-mini (D) and LARI-1.0 (E). **(F)** Fully assembled dual-probe LARI-1.0 and LARI-mini with ground cable and M1 screw weighed ~1.8 and ~1.0 grams respectively (without the headstage). Adding the headstage weight (5-6 grams) the total weight of the dual (~2.4 g) and single (~1.6) is around 10% and 7% of the average weight of adult C57BL6 mice.

To test the implant systems, animals underwent between 18 and more than 60 recording sessions across several weeks to months. Each mouse participated in repeated sessions spaced 5–7 days apart across three metabolic conditions: fed state, 24-hour fasted state, and high-fat-diet (60% HFD)-induced obesity state. Due to occasional mortality or implant breakage caused by user handling errors, not all animals completed all metabolic phases; however, roughly 80% (n = 4) completed all three. Using LARI-mini, we recorded from retrosplenial cortex (RSP), hippocampal CA1, and lateral hypothalamus (LHA). With dual-probe LARI-1.0, we recorded from the same regions plus the anterior cingulate area (ACA).

LARI-1.0 can be used with either a single or two Neuropixels 2.0 probes (**Figure 2A**). LARI-mini is designed specifically for single-probe use and reduces implant weight by an additional ~20% compared to LARI 1.0 (**Figure 2C and F**). LARI-1.0 supports adjustable relative depth and lateral spacing between two probes (**Figure 2B**), enabling focal targeting of cortical and subcortical regions in one or both hemispheres. Figure 2B (*right*) shows the schematic of how LARI-1.0 allows independent adjustment of the two probes to selectively target ACA and RSP/LHA—two narrow cortical structures positioned near the midline—without needing to rotate the implant or encroaching on adjacent structures. This flexibility is a key feature of the LARI-1.0 design that expands capabilities to record from different brain regions compared to other solutions.

LARI-1.0 includes three components: the skull mount (SM), a recoverable internal module (IM), and a protective cap, whereas LARI-mini only consists of SM and IM. The LARI-1 IM contains a central plane with vertical and horizontal grooves spaced 0.5 mm apart for fine depth and lateral adjustments (**Figure 2B**). Probes are affixed to opposite sides of this plane. The flex cable and headstage are housed inside the IM, which slides into the SM and is secured with M1.2 screws. The cap attaches to the IM using screws or glue and contains a threaded brass insert for stereotaxic attachment. The SM consists of a channel system (**Figure 2A**, *far right*) with a housing surrounding the probes that attaches to the skull. This channel allows for further fine adjustments of effective shank length after fixing the probe depth in the IM. Though calculations of required probe depth pre-assembly of probes in the IM can be made close to the desired accuracy, in our case in some cases we found that it is always better to leave some room for final adjustments right before insertion to ensure that desired length of shank (insertion depth in DV + depth of dura + ~ 1mm of space between the SM tip and skull) can be attained. We tested SM’s (both mini and LARI-1) with a fully enclosed design for covering the probes and also a small opening in the back (side facing the surgeon), in our experience we found it more practical to leave an opening in the back (**Figure 2D and 2E**) that allows viewing the probe shanks for any deformations or kinks during insertion. This opening is covered towards the end of surgical implantation (described in Methods and Supplementary material).

LARI-mini includes a skull mount and an internal module with an integrated stereotax sleeve for stereotax rod (**Figure 2C**, *far right*). In both LARI-1.0 and LARI-mini, the ground wire is internally routed to minimize electromagnetic noise (**Figure 2A, 2C**). Similar to LARI-1, SM in the LARI-mini also permits adjustment of the effective shank depth via a sliding channel interface secured by M1 screws (**Figure 2C**). In all uses, a single M1 screw per side was sufficient to secure the SM-to-IM connection.

The fully assembled LARI-1.0 implant (without headstage) including SM, IM, two Neuropixels-2.0 probes, cap, superglue, two M1 screws, 4 brass inserts, and ground cable attached to an M1 screw weighs approximately 1.8 grams (**Figure 2G,F**), making it one of the lightest dual-probe systems for chronic mouse recordings. The 3D-printed implant components alone weighed ~1.0–1.1 grams depending on printing material. A fully assembled LARI-mini without headstage weighs ~0.9–1.0 grams, with printed components weighing 0.58–0.69 grams (**Figure 2F**). Including the 0.5-gram headstage, total implant mass was ~2.3 grams for LARI-1.0 and ~1.5 grams for LARI-mini, approximately 10% and 7.5% of average male mouse body weight, respectively. This is intended to reduce the burden on the mice to help enable natural behavior. In summary, our design provides a flexible, lightweight solution for chronic *in vivo* recordings from varied brain regions.

### Implant Stability

To evaluate the performance of our system for capturing chronic recordings from foraging mice, we first evaluated implant stability over time. Based on observations of foraging behavior, we hypothesized that the most significant mechanical stress on the implant arises from neck, head, and craniofacial muscle movements during high-movement behaviors such as digging, chewing, and grooming. To examine whether these forces generated artifacts, we isolated digging and feeding bouts from behavioral data and extracted corresponding spike rasters from the LARI-1.0 dual probe recorded mice. We found no consistent high-density artifact spikes across units or depths associated with digging or feeding (**Figure 3A**). Examination of spike activity across the full range of mouse velocities also revealed no systematic correlation between high-velocity bouts and spike artifacts (**Figure 3B**). Occasional artifacts were observed, but they were not consistent across fast movements and therefore unlikely to be motion related. Similar results were observed for LARI-mini. Thus, both probe systems yielded stable recordings with few overt artifacts.

**Figure 3.**
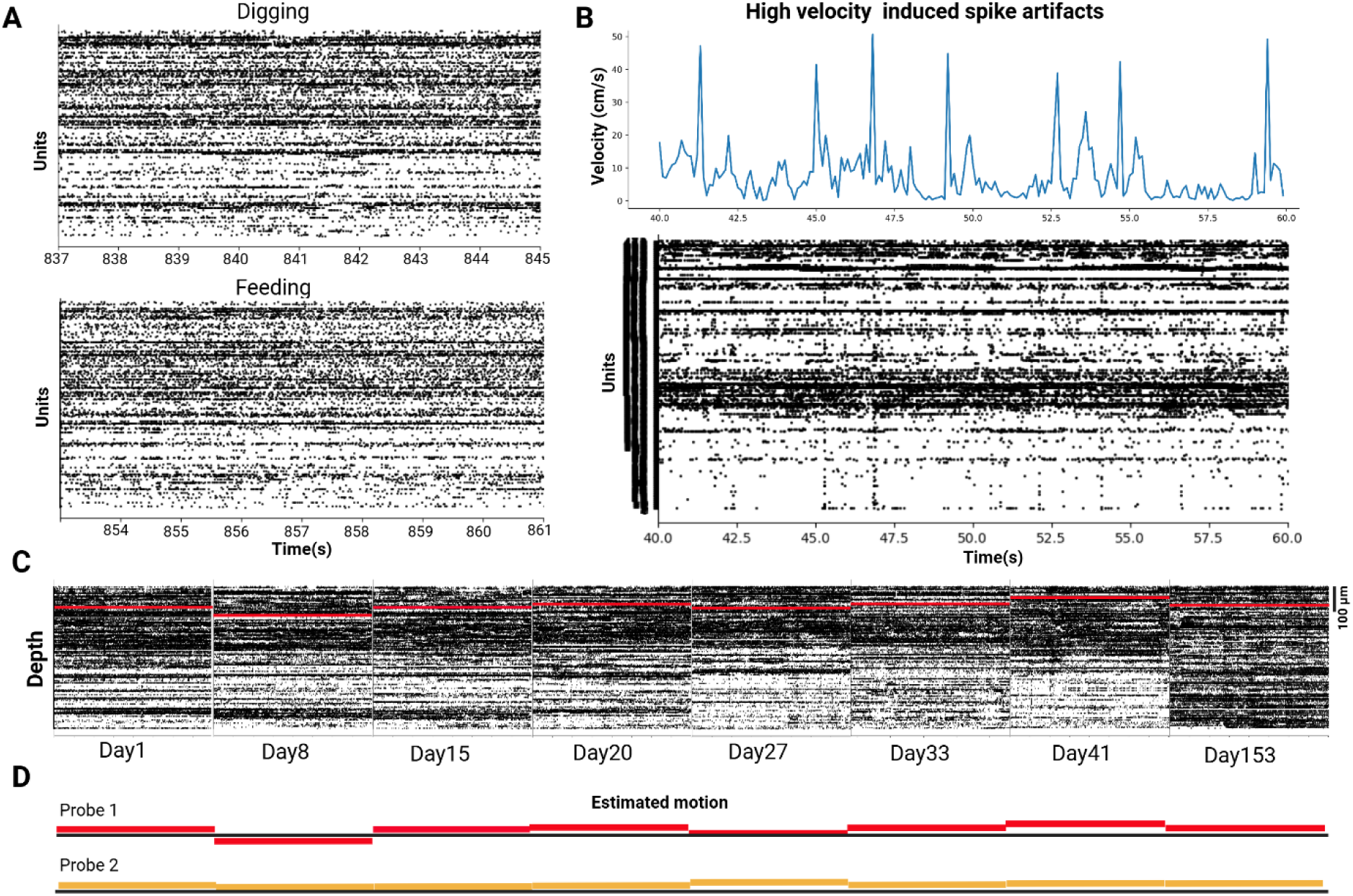
Evaluation of LARI-1.0 dual implant stability and motion artifacts. **(A)** Raster plot of all units from a recording during a digging and feeding window. Naturalistic behaviors such as digging involve intense limb movements that can introduce vibrations and artifacts in the recordings and jaw movements while feeding is known to also introduce licking-like artifacts. In our case, we did not observe the sharp high-density artifacts usually observed in freely moving *in vivo* recordings, demonstrating that our platform captures high quality data during key natural behaviors. **(B)** We encountered some spikes that seem to coincide with high-velocity movement bouts and are suspicious for being technical artifacts. However, they were not consistent across all high-velocity events, therefore ruling them out as entirely originating from animal movement. This suggests that many reflect bona fide biological neural activity. **(C)** Probe drift/motion estimated by concatenating the same segment of recordings across all sessions. The results show minimal drift and indicate stable recordings across sessions. **(D)** With dual probe LARI-1.0, probe 1 targeting the anterior cingulate cortex (ACA) shows relatively more motion than probe 2 targeting retrosplenial cortex (RSP) and lateral hypothalamus (LHA). Our estimated motions (both single and dual probes) were similar to, or in some cases lower than other published studies (see main text).

We next assessed long-term probe drift/motion in dual-probe LARI-1.0 implants over more than 180 days (data shown to 153 days; **Figure 3C and 3D**) using approaches used by other studies^22,31^. Drift magnitude was similar to that found by these studies ^22,31^. **Figure 4A and B** show probe motion across days/sessions of single Neuropixels-2.0 probe implanted with LARI-mini. LARI-mini implants, had almost half the amount of probe motion reported by recording systems in other studies (median displacement of 1.93 ± 2.83 µm for one implant shown in **Figure 4A**). Across all LARI-mini implantations (n=5) the median displacement was contained within 1.5 to 2.35 ± 3 µm. Similar to other studies^17,22^, we also encountered probe motion to be highest in the first month after implantation (**Figure 4A** and **4B**, *top*), which has been suggested to relate to surgical recovery and easily correctable with registration algorithms ^17,22^. Figure 4B shows data from a reused probe, the probe first used in the mouse shown in figure 4A. Figure 4A and 4B (*middle* and *bottom*) shows a depth plot of matched units across all sessions from LHA and RSP. Overall, the LARI-mini exhibits minimal drift and yields stable recordings that allowed us to track units over time.

**Figure 4.**
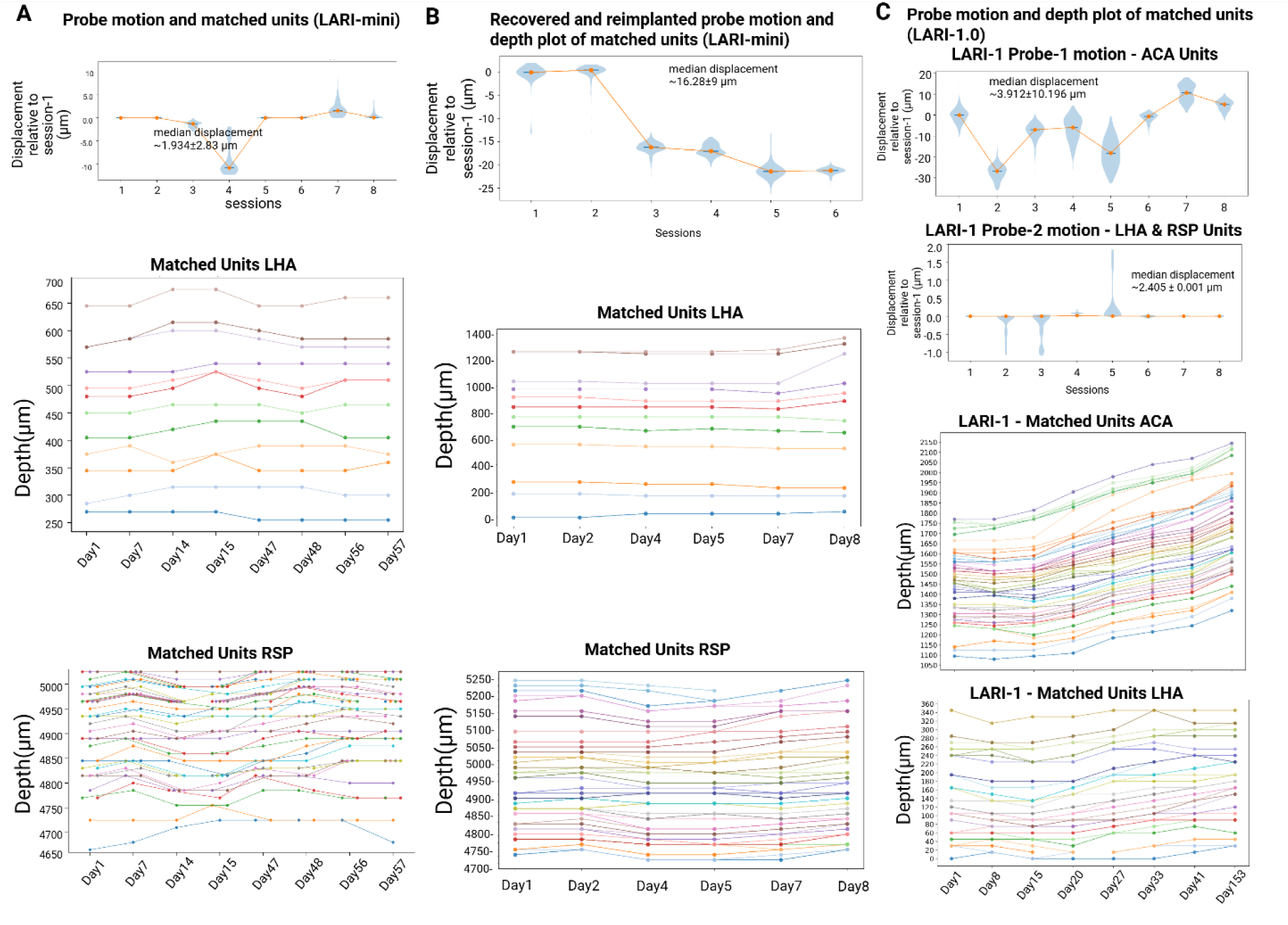
Stable unit recordings are observed for chronic LARI-mini and LARI-1.0 recordings. **(A)** Global probe motion (top) and depth plot of matched LHA (middle) and RSP (bottom) units across days recorded from a single Neuropixels-2.0 probe implanted with LARI-mini. We observed probe drift/motion similar to reported by study with permanently cemented probes^17^. Our median probe displacement (shown here for one implanted probe ~1.934 ± 2.83 µm) calculated with an established method^31^ was equally comparable and in some cases smaller than what was reported by other studies^17,22^. **(B)** Probe motion and depth plot of matched units from a recovered and reused probe implanted with LARI-mini. **(C)** Motion of probe1 and 2 in the dual-probe LARI-1.0 (*top* and *middle*) and depth plot of matched units from ACA (probe1) and LHA recorded (probe2). Similar to other reports^22^, we observed most of the probe motion in the first month after implantation, which seem to stabilize after (as seen in *top* plots in A, B, and C). ACA matched units (C, second from *bottom*) show higher depth changes than the observed global probe motion, which could be due to inherent bias in manual sorting to match units and/or intercellular motion that cannot be explained entirely by probe motion alone.

For LARI-1.0 dual probe implants, the probe targeting the cortical region (ACA targeting probe1 - median displacement 3.912±10.196 µm, **Figure 4C** *top*) showed increased motion compared to the hypothalamic units (LHA targeting probe2 - median displacement 2.405±0.001 µm, **Figure 4C** *middle*). This is comparable with studies that found that cortical units show more motion (4.9±7.6 µm displacement) through sessions compared to units from ventral brain areas such as thalamus (1.6±8.2 µm displacement) ^22^. The regional differences could be due to immune responses during the first month after probe implantation, in our case, we also suspect the motion in probe1 may arise from age-dependent craniofacial growth or increased muscle bulk—exacerbated by the mouse’s 30–35% weight gain during the 60-day high-fat-diet period. Then we examined depth changes of manually matched units across sessions. We separated units from the two brain regions to see if motion affects different brain regions differentially and if dorsal (RSP) and ventral (LHA) regions have any unique motion characteristics. **Figure 4C** (*bottom*) shows the depth of matched units in the ACA across sessions from probe1 and in the LHA from probe2. The overall pattern of depth changes of matched units was slightly higher for cortical regions (**Figure 4C**, compare ACA to LHA). This is similar to the pattern observed in single probe implantations using LARI-mini (**Figure 4A** and **4B**). The matched unit depth changes are a bit higher than the estimated full-probe motion numbers, which is expected because the latter is a cumulative approach and individual units/neurons can have depth changes different than the observed direction of cumulative probe motion, owing to discrepancies due to manual curation as well immune and cellular factors that may affect units differentially in different depths.

Overall, both LARI-1.0 and LARI-mini remained highly stable throughout the study and produced results comparable or better than prior studies, successfully isolating the probes from motion and vibration-induced forces arising during naturalistic behaviors. Of the four implant breakages encountered, three occurred due to user error during handling or probe cleaning. The only breakage occurring on an implanted mouse resulted few hours post-implantation due to insufficient bonding between skull and dental cement resulting from inadequate skull surface preparation. All failures were resolved with prolonged surgical training with dummy probes.

### Spike Quality

To evaluate the data quality collected with our probe design and naturalistic foraging paradigm, we used a conservative combinatorial curation approach, integrating manual inspection (Phy, https://github.com/cortex-lab/phy) with automatic curation^32^ (SpikeInterface), and included only units that passed multiple spike-quality metrics including waveform shape, spike amplitude, interspike interval (ISI), spike pattern, unit location, and depth. Because template-based unit matching in SpikeInterface occasionally misassigned units, all automatically identified units were manually verified.

Figures 5A and 5B show average spike waveform and interspike intervals of three different units recorded with probes implanted with LARI-mini and LARI-1.0. Average waveform shapes and ISI histograms remained stable across sessions for most units (Figures 5A**, B**). Some ISI and amplitude variability were expected due to the free-choice nature of the task, in which reward location, physiological states, and spontaneous behavioral sequences are varied. Figure 5C and 5D show representative traces from probes targeting RSP, LHA and ACA. All single-probe LARI-mini recordings simultaneously targeted RSP, hippocampus (not shown), and LHA, whereas dual probe LARI-1.0 targeted RSP, LHA and ACA.

**Figure 5.**
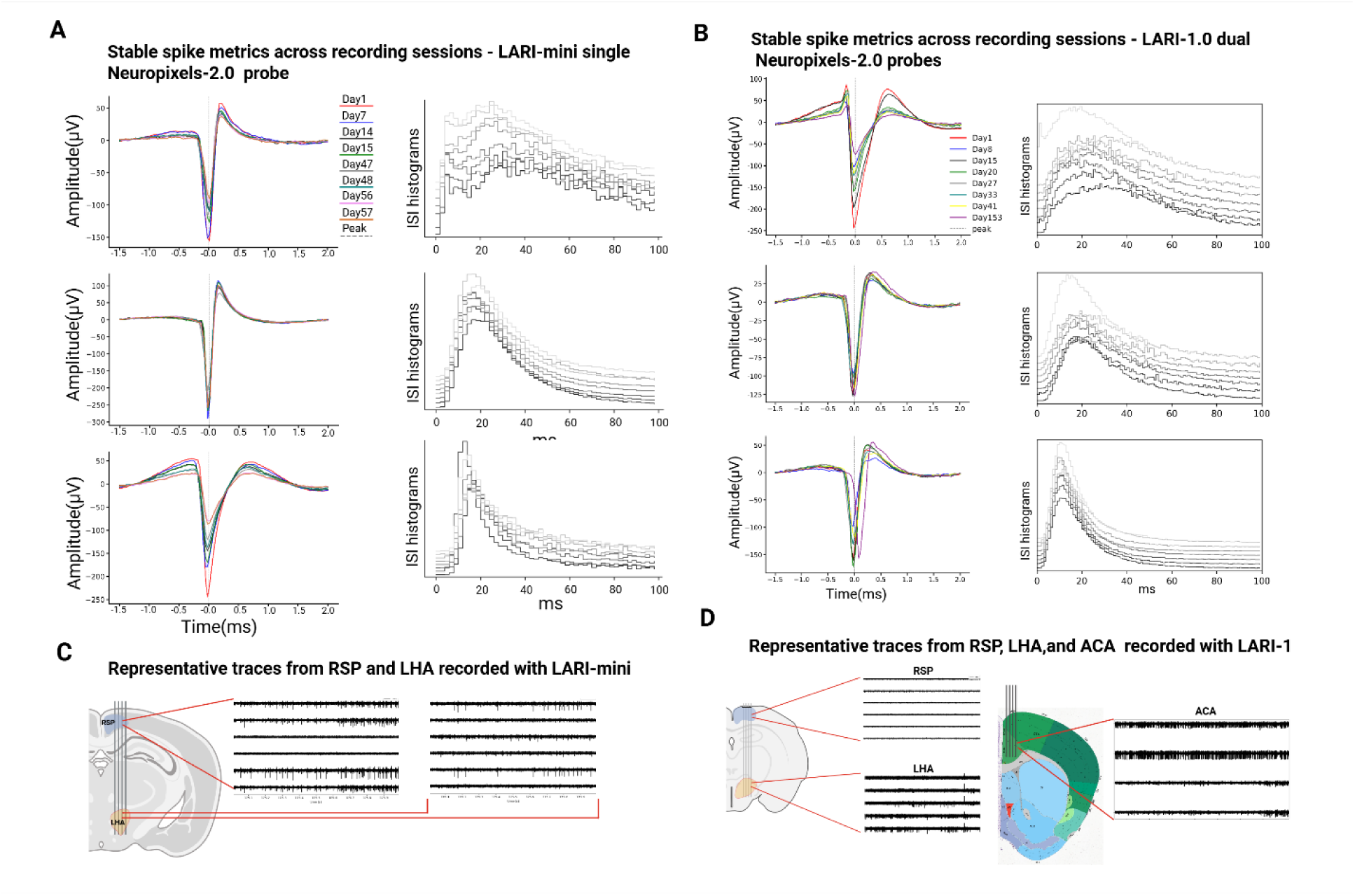
Spike quality metrics of units recorded with probes implanted with LARI-mini and LARI-1.0. **(A)** Average waveform and ISI of three units matched across sessions recorded from a single Neuropixel-2.0 probe implanted with LARi-mini. **(B)** Representative traces from channels located in RSP and LHA. In general, units in RSP had higher firing rates and total number of spikes. **(C)** Average waveform and ISI of three units matched across sessions recorded from dual Neuropixel-2.0 probes implanted with LARI-1.0. **(D)** Representative traces from channels located in RSP and LHA (Probe 2) and ACA (probe 1). We found more units in ACA when compared to both RSP and LHA.

Signal-to-noise ratios were consistently in the 5–7-fold range, indicating that robust neural activity signals are captured in the data for the LARI-mini (Figure 6A, left), recovered LARI-mini (Figure 6B, left), and LARI-1.0 systems (Figure 6C, left). Mice underwent major metabolic shifts (fed, fasted, and HFD), which likely contributed to changes in firing patterns across different days. Good unit yields also remained high across recording sessions with mice showing last session unit yields similar to the first day of recording (Figure 6A-C, right). Overall, unit yields remained within ~20% of the unit yield on the first recording day for all animals. We did not quantify amplitude changes across sessions because amplitude changes were inconsistent across units partly owing to the biological variability of the expressed natural foraging behavior and could not be attributed solely to technical factors, such as drift or signal deterioration.

**Figure 6.**
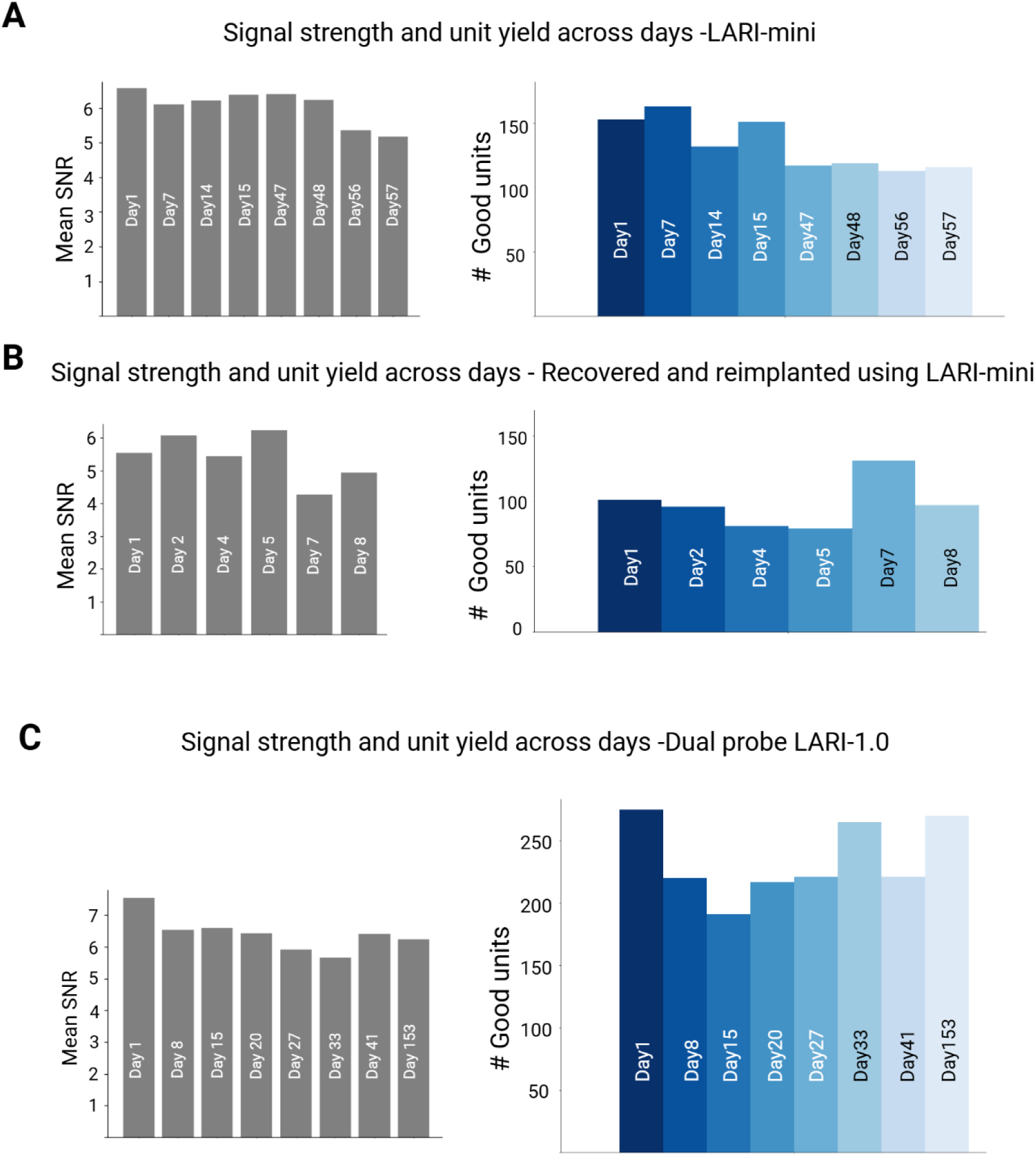
Signal-to-noise ratio (SNR) and Kilosort labeled ‘good’ single unit yields across sessions and implant systems. **(A)** Mean SNR and number of good units (combined LHA and RSP) in a single probe implanted with LARI-mini. **(B)** Mean SNR and good unit yield from a mouse implanted with a reused probe using LARI-mini. **(C)** Mean SNR and good unit yield from two probes (RSP, LHA, and ACA) implanted with LARI-1.0.

### Unit yield

We investigated capabilities to track individual units across recording sessions in our naturalistic foraging task using LARI-mini and LARI-1.0 recordings. The behavior in this task is spontaneous and variable such that individual units are expected to show more intrinsic variability and response diversity across sessions compared to constrained operant conditioning tasks. Nonetheless, we found that the number of Kilosort-3^33^ labeled “good” single-units did not exhibit consistent declines or significant changes for LARI-mini or LARI-1.0 recordings (*left* in Figures 6A-C). Units may disappear or newly emerge across days due to immune responses^34–36^ at the tissue–probe interface, neuronal loss, or behavior-dependent shifts in firing patterns. In an initial analysis of unit capture, we used SpikeInterface-based automated curation and found a high proportion of matched units (~90%) across sessions. Next, we adopted a conservative matching strategy to identify the same units across sessions with high confidence by manually inspecting spike waveform, firing rate, and temporal structure after SpikeInterface-based automated curation. This approach discarded a substantial number of good quality single units with 60% fewer units retained on the final recording day compared to the first recording day. The number of high confidence matched units across sessions between different animals remained within a range of 50±5 for RSP and 10±4 for LHA for single-probe LARI-mini implants, and 55±4 (ACA) and 14±2 (LHA) for LARI-1.0 dual-probe implants. Thus, our systems successfully tracked the activity of many units matched with high confidence across sessions from natural foraging mice, enabling comparative analyses over time. We expect that unit yield would be higher in more constrained experimental paradigms, where neural activity varies less across sessions and unit matching is therefore more straightforward. Similar improvements would also be expected in acute, head-fixed preparations with tightly controlled task structure.

## Discussion

Neuroscience has traditionally emphasized tightly controlled tasks that isolate single processes, whereas naturalistic studies seek to understand how those processes operate within the dynamic, spontaneous, multidimensional flow of real life. By expanding beyond simplified, trial-based designs or head-fixed paradigms, more naturalistic behavioral paradigms can reveal novel mechanisms, components, and emergent properties—such as patterns, strategies, and cross-system coordination—that remain largely unexplored. Neuropixels high density recordings across multiple brain regions in foraging mice offer the potential to discover new insights into the neural computations underlying motivational, learning and memory, homeostatic, cognitive, and decision processes. Here, to advance the application of neuropixels technology for naturalistic behaviors, like foraging, we developed and demonstrated the LARI-mini and LARI-1.0 implant fixtures compatible with both single- and dual-Neuropixels 2.0 probes. The system enables independent adjustment of probe depth and lateral spacing, facilitates probe recovery and reuse, and provides sufficient mechanical stability for long-term single-unit recordings from multiple mouse brain regions during naturalistic foraging in a feature-rich environment over weeks to months.

Using these implants, we recorded from multiple regions within the same hemisphere, including the ACA, LHA, and RSP. A key feature of our design is the ability to selectively target the brain regions under study while minimizing probe traversal through non-targeted areas. This feature reduces unnecessary shank insertion into adjacent brain regions and helps avoid potential confounds arising from off-target tissue disruption.

Several existing implant designs are lightweight and enable independent probe insertion angles, such as the Apollo implant^25^ and the *indie* system^22^. However, these designs lack the adaptability afforded by our approach, particularly with respect to selectively targeting brain regions through independent adjustment of probe depth and lateral spacing within the internal module (IM), as well as additional depth adjustment via the skull mount (SM). This flexibility allows precise targeting along both the anteroposterior (AP) and dorsoventral (DV) axes while minimizing probe traversal through non-targeted brain regions. Using this approach, we were able to selectively access regions of interest while leaving surrounding tissue undisturbed. Both inter-session and intra-session brain–probe motion were reduced relative to reports using permanently cemented probes^17^, while yielding a similar number of well-isolated single units. Finally, our naturalistic freely foraging study design also included three different metabolic phases (Standard chow ‘Fed’, 24-hour ‘Fasted’, and 60% high-fat diet ‘Fed’) and in spite of these metabolic interventions, resulting in significant weight loss/gain and behavioral changes, unit quality metrics and unit yield remained stable and reliable across sessions.

### Caveats

Although the dual-probe LARI-1.0 design supports implantation of two Neuropixels probes within a single hemisphere or across both hemispheres, the current configuration does not readily accommodate simultaneous targeting of anatomically distant regions, such as the cerebellum and frontal cortex. Targeting such regions would require user-driven customization of the implant, including increasing the thickness of the central plane within the internal module (IM) and corresponding modifications to the skull mount (SM). In addition, probe attachment within the IM can be technically challenging for some users, as the flex cables must be routed through the headstage housing prior to probe fixation. To address these limitations, future iterations of the LARI platform will focus on expanding anatomical targeting flexibility to support both nearby and widely separated brain regions, as well as developing a more user-friendly probe attachment and alignment interface.

### Future directions

In addition to the future directions outlined above, we plan to extend the LARI platform to larger animals, such as rats, and to increase the number of probes that can be simultaneously implanted within a single fixture (up to six probes in rats and four probes in mice). This expansion can be achieved primarily by scaling the existing components, without requiring major changes to the underlying design. We also plan to incorporate customizable adjustment of inter-probe spacing and insertion angles within a single fixture to enable simultaneous targeting of both anatomically proximal and distant brain regions. Further reductions in implant size and weight are also planned. For example, the combined mass of threaded brass inserts and screws currently accounts for approximately 0.3–0.4 g of the implant weight and could be substantially reduced by redesigning the cap to eliminate brass inserts in favor of a threaded stereotax rod that directly interfaces with the cap. In addition, we found that the wall thickness of several components can be reduced by ~0.2 mm when printed using Tough 2000 resin on Formlabs Form 4 printers, potentially yielding additional weight savings without compromising mechanical strength or flexibility. Finally, we plan to make updated and improved design files publicly available through a GitHub repository and to provide free consultation to support customization and adoption of the LARI system by the research community.

Our work has been possible due to the miniaturization and probe development efforts done by others^14,17^ The field is under active development with newer and lighter probe and headstage models that will be publicly available in the upcoming years. We will make changes in our design accordingly to accommodate any changes in the newer models and offer flexible adjustments to maximize accessibility, selective targeting, and reuse of the probes.

## Materials and Methods

### Implant Design and 3D Printing

All implant designs are available in the github repository https://github.com/gregglab-org/LARI-implant-. More design file formats will be added in the near future and any future iterations of the design will be updated in the repository accordingly.

We used multiple 3D printing materials for fabrication of both LARI-1.0 and LARI-mini implants, including Nylon PA12 and Formlabs Tough 2000 resin. PA12 components were produced using either selective laser sintering (SLS) or multi jet fusion (MJF), while Tough 2000 parts were fabricated using Formlabs stereolithography (SLA). In general, SLS-printed PA12 implants were consistently lighter (by ~15–20%) than MJF-PA12 and Tough 2000 parts, but required greater wall thickness to maintain structural integrity. In contrast, Tough 2000 offered higher flexibility and tensile strength at the cost of a 15–20% increase in weight.

All implant components were designed using Autodesk Fusion 360, and STL files were generated for printing. Tough 2000 fixtures were printed using Formlabs Form 3+ printers at the Center for Medical Innovation (CMI) Prototyping Services at the University of Utah School of Medicine. Nylon PA12 SLS and MJF printing was outsourced to third-party services (Shapeways and Xometry). All SLA-printed parts were post-processed according to Formlabs-recommended washing, drying, and curing protocols. All fixtures were printed in a vertical orientation, and we found that orienting the anterior end (skull facing) of both the internal module (IM) and skull mount (SM) toward the build platform yielded the best dimensional accuracy and print consistency.

### Probe Assembly

Two to four M1 threaded brass inserts (McMaster-Carr) for plastics (Figure 2) were heat-tapped into each IM and two into the protective cap. These inserts enabled fastening between the external module (EM) and IM, as well as cap-to-stereotax connections. A Neuropixels 2.0 probe was first carefully slid into the downward-facing side of the IM central plane and secured with cyanoacrylate glue at the desired depth and lateral position. A second probe was then positioned and secured on the upward-facing side of the central plane. Flex cables from both probes exited through the headstage housing at the rear of the IM.

An Ag/AgCl ground wire was looped through the ground pads on the flex cables of both probes and soldered carefully to avoid overheating or damaging the flex cables. The ground wire was routed internally through the headstage housing and the EM channel. The EM was then slid onto the IM and fastened using M1 screws on both sides of the EM–IM interface. The protective cap was placed onto the upper flanges of the IM and secured using either M1 screws or cyanoacrylate glue. Screws were tightened only until secure, as excessive torque during tightening or loosening could induce probe shank breakage during recovery. A single M1 ground screw was soldered to the ground wire.

### Trajectory Planning and Visualization

Probe trajectories were planned using the Pinpoint trajectory planner^37^. The graphical user interface was used to determine anteroposterior (AP), mediolateral (ML), and dorsoventral (DV) coordinates, as well as relative spacing and depth between probes. Because our study required simultaneous targeting of ACA, RSP, and LHA, we used Pinpoint to identify appropriate AP spacing that would allow coverage of these regions. These constraints informed the design width of the central plane in the LARI-1.0 IM. The Pinpoint API combined with spikeGLX was also used to configure probe channel banks across the four shanks to maximize coverage of regions of interest.

### Surgical Implantation

All animal procedures followed NIH guidelines and were approved by the Institutional Animal Care and Use Committee (IACUC) at the University of Utah. A total of seven C57BL/6 mice (five males and two females) were used in this study.

A detailed surgical protocol is provided in the GitHub repository and supplementary materials. We followed protocols similar to other studies^18,38^ and modified them accordingly for our design and study. Male and female mice aged 8–10 weeks were anesthetized using 3% isoflurane and mounted on a Stoelting stereotaxic frame equipped with a mouse-specific nose cone. Adequate anesthesia was confirmed using a twitch reflex test, after which lidocaine (2–4 mg/kg) was administered subcutaneously over the skull. After a 5–8 min diffusion period, a midline incision was made to expose the skull. The skin edges were secured to the skull using cyanoacrylate tissue adhesive (Vetbond, World Precision Instruments) to maintain exposure.

After stereotaxic alignment, craniotomy locations were marked by lightly etching the skull with a dental drill. For single-probe LARI-mini implants, three coordinate sets were used to target the AP extent of RSP and LHA (AP: −1.1, −1.5, or −2.0 mm; ML: 0.74 mm; DV: 6.1 mm). For dual-probe LARI-1.0 implants, probe-1 targeted ACA (AP: 0.99 mm; ML: 1.16 mm; DV: 3.27 mm), while probe-2 targeted RSP and LHA (AP: −2.0 mm; ML: 0.74 mm; DV: 6.1 mm; Figure 6E). AP and ML reference lines were etched to aid alignment during drilling. The dura was removed to facilitate probe insertion. A contralateral burr hole was drilled for placement of the ground screw.

The skull surface was etched with a scalpel blade, dried, and a ring of dental composite resin (Charisma, Kulzer; Net32 cat. no. 66000085) was created encircling the craniotomies to sustain a well of saline during insertion to keep the exposed brain moistened and facilitate insertion of probes (Figure 2C). The fully assembled implant, including headstage, was attached to the stereotax rod, and probes were inserted at 10 μm/s using an automated micromanipulator (World Precision Instruments). Prior to insertion shanks were sterilized with isopropanol and labeled with CM-DiI (Thermo Fisher Scientific) for post hoc histological verification.

We adjusted our shank depth and the SM connection to IM such that on complete dorso-ventral insertion the EM lightly contacts the skull or remains within <1 mm of the surface. The space inside the IM were filled with biocompatible silicon sealant (Kwik-cast, WPI) to ensure lower probe movement. The opening on the back of IM in mini and LARI-1.0 implants were covered with a small copper tape adhesive (or thin plastic) and later covered with dental cement. Premixed dental resin (3M RelyX Unicem Automix) was applied at the SM–skull interface, and the ground screw was secured with dental resin. After curing, dental cement (Metabond; mixed cold at 2 scoops powder, 4 drops liquid, and 1 drop catalyst) was applied to secure the implant. Care was taken to fully encapsulate exposed ground wire and avoid contact between cement and surrounding skin or fur. After 7–10 min of curing, stereotax screws were removed and the rod was slowly retracted at 15–20 μm/s.

Carprofen (5 mg/kg) was administered subcutaneously prior to recovery. Mice were monitored in a heated recovery cage and returned to housing once normal behavior resumed. Carprofen treatment continued for 2 additional days, and recordings began no earlier than 10 days post-surgery.

### Simultaneous Spike Acquisition and Naturalistic Foraging Behavior

Mice underwent naturalistic foraging sessions consisting of three phases. The first 15 min were restricted to the home cage, followed by a 30 min exploration phase initiated by automatic opening of the door, with visible seeds placed in one pot. After a 2-hour intermission, a 30 min foraging phase was conducted with seeds buried beneath one pot (distinct from the exploration phase). Behavioral recordings were not collected during the intermission.

Behavior was recorded using EthoVision XT (Noldus) with GigE Basler acA1300-60gmNIR cameras, and movement was tracked using deep-learning-based three-point tracking. Neural data were acquired simultaneously using SpikeGLX. Behavioral events, camera frame timing, and door-opening signals were recorded as TTL inputs and synchronized post hoc using TPrime.

### Spike Sorting and Curation

Raw neural data were preprocessed using a 300–9,000 Hz third-order Butterworth bandpass filter, followed by phase-shift correction and common median subtraction. Spike sorting was performed using Kilosort 3 with default parameters. Preprocessing, sorting, and postprocessing were conducted using the SpikeInterface framework and custom Python scripts for analysis and visualization.

Both automated (SpikeInterface) and manual (Phy) curation were used. Kilosort-3 “good” labels were not relied upon exclusively due to known misclassification issues. All units were manually evaluated based on waveform shape, amplitude, ISI distributions, and cross-correlograms. Units were matched across sessions using SpikeInterface’s template comparison module; all automated matches were manually verified, and incorrect matches were excluded.

### Probe Recovery, Cleaning, and Reimplantation

For probe recovery, mice were anesthetized with 3% isoflurane and mounted in a stereotaxic frame with the implant aligned near 90° relative to the horizontal plane. The stereotax rod was secured to the cap (LARI-1.0) or integrated sleeve (LARI-mini) using M1 or M1.4 screws. The ground wire was exposed by gently removing dental cement with a dental drill and then severed. M1 screws connecting the SM and IM were removed, and the IM was slowly retracted at 10–15 μm/s until the probe shanks were fully exposed and the SM separated.

Probes were cleaned without removal from the IM by soaking in warm (<37°C) Tergazyme for 1–2 hours or sonicating at room temperature for 30 min, followed by sterilization in isopropanol for 10–20 min. Shanks were inspected under a stereo microscope and cleaning was repeated if necessary. Reassembly required only resoldering the ground screw and reattaching the SM to the IM. Reimplantation followed the same procedure as the initial surgery.

## Acknowledgements

We thank all members of the Gregg Lab for helpful discussions. D. Joseph from center of medical innovation (CMI) at the University of Utah school of medicine for his patience with countless iterations and prints, prototyping and troubleshooting print issues.

## References

1. Huk A, Bonnen K, He BJ. Beyond Trial-Based Paradigms: Continuous Behavior, Ongoing Neural Activity, and Natural Stimuli. J Neurosci. 2018;38(35):7551–7558. PMCID: PMC6113904

2. Nathan R, Monk CT, Arlinghaus R, Adam T, Alós J, Assaf M, Baktoft H, Beardsworth CE, Bertram MG, Bijleveld AI, Brodin T, Brooks JL, Campos-Candela A, Cooke SJ, Gjelland KØ, Gupte PR, Harel R, Hellström G, Jeltsch F, Killen SS, Klefoth T, Langrock R, Lennox RJ, Lourie E, Madden JR, Orchan Y, Pauwels IS, Říha M, Roeleke M, Schlägel UE, Shohami D, Signer J, Toledo S, Vilk O, Westrelin S, Whiteside MA, Jarić I. Big-data approaches lead to an increased understanding of the ecology of animal movement. Science. 2022;375(6582):eabg1780. PMID: 35175823

3. Cisek P, Green AM. Toward a neuroscience of natural behavior. Curr Opin Neurobiol. 2024;86:102859. PMID: 38583263

4. Dennis EJ, Hady AE, Michaiel A, Clemens A, Tervo DRG, Voigts J, Datta SR. Systems Neuroscience of Natural Behaviors in Rodents. J Neurosci. 2020;41(5):911–919. PMID: 33443081

5. Kennedy A. The what, how, and why of naturalistic behavior. Curr Opin Neurobiol. 2022;74:102549. PMCID: PMC9273162

6. Budaev S, Jørgensen C, Mangel M, Eliassen S, Giske J. Decision-Making From the Animal Perspective: Bridging Ecology and Subjective Cognition. Frontiers Ecol Evol. 2019;7:164.

7. Huang WC, Ferris E, Cheng T, Hörndli CS, Gleason K, Tamminga C, Wagner JD, Boucher KM, Christian JL, Gregg C. Diverse Non-genetic, Allele-Specific Expression Effects Shape Genetic Architecture at the Cellular Level in the Mammalian Brain. Neuron. 2017;93(5):1094–1109.e7. PMID: 28238550

8. Mobbs D, Trimmer PC, Blumstein DT, Dayan P. Foraging for foundations in decision neuroscience: insights from ethology. Nat Rev Neurosci. 2018;19(7):419–427. PMID: 29752468

9. Grima LL, Haberkern H, Mohanta R, Morimoto MM, Rajagopalan AE, Scholey EV. Foraging as an ethological framework for neuroscience. Trends Neurosci. 2025;48(11):877–890. PMCID: PMC12693718

10. Hörndli CNS, Wong E, Ferris E, Bennett K, Steinwand S, Rhodes AN, Fletcher PT, Gregg C. Complex Economic Behavior Patterns Are Constructed from Finite, Genetically Controlled Modules of Behavior. Cell Rep [Internet]. 2019;28(7):1814–1829.e6. Available from: 10.1016/j.celrep.2019.07.038 PMCID: PMC7476553

11. Bonthuis PJ, Steinwand S, Hörndli CNS, Emery J, Huang WC, Kravitz S, Ferris E, Gregg C. Noncanonical genomic imprinting in the monoamine system determines naturalistic foraging and brain-adrenal axis functions. Cell Rep. 2022;38(10):110500. PMCID: PMC9128000

12. Ravens A, Stacher-Hörndli CN, Emery J, Steinwand S, Shepherd JD, Gregg C. Arc regulates a second-guessing cognitive bias during naturalistic foraging through effects on discrete behavior modules. iScience. 2023;26(5):106761. PMCID: PMC10196573

13. Steinwand S, Hörndli CS, Ferris E, Emery J, Murcia JDG, Rodriguez AC, Spotswood RJ, Chaix A, Thomas A, Davey C, Gregg C. Conserved noncoding cis elements associated with hibernation modulate metabolic and behavioral adaptations in mice. Science. 2025;389(6759):501–507. PMCID: PMC12403242

14. Jun JJ, Steinmetz NA, Siegle JH, Denman DJ, Bauza M, Barbarits B, Lee AK, Anastassiou CA, Andrei A, Aydın Ç, Barbic M, Blanche TJ, Bonin V, Couto J, Dutta B, Gratiy SL, Gutnisky DA, Häusser M, Karsh B, Ledochowitsch P, Lopez CM, Mitelut C, Musa S, Okun M, Pachitariu M, Putzeys J, Rich PD, Rossant C, Sun W lung, Svoboda K, Carandini M, Harris KD, Koch C, O’Keefe J, Harris TD. Fully integrated silicon probes for high-density recording of neural activity. Nature. 2017;551(7679):232–236. PMCID: PMC5955206

15. Chung JE, Sellers KK, Leonard MK, Gwilliams L, Xu D, Dougherty ME, Kharazia V, Metzger SL, Welkenhuysen M, Dutta B, Chang EF. High-density single-unit human cortical recordings using the Neuropixels probe. Neuron. 2022; PMID: 35679860

16. Paulk AC, Kfir Y, Khanna AR, Mustroph ML, Trautmann EM, Soper DJ, Stavisky SD, Welkenhuysen M, Dutta B, Shenoy KV, Hochberg LR, Richardson RM, Williams ZM, Cash SS. Large-scale neural recordings with single neuron resolution using Neuropixels probes in human cortex. Nat Neurosci. 2022;25(2):252–263. PMID: 35102333

17. Steinmetz NA, Aydin C, Lebedeva A, Okun M, Pachitariu M, Bauza M, Beau M, Bhagat J, Böhm C, Broux M, Chen S, Colonell J, Gardner RJ, Karsh B, Kloosterman F, Kostadinov D, Mora-Lopez C, O’Callaghan J, Park J, Putzeys J, Sauerbrei B, Daal RJJ van, Vollan AZ, Wang S, Welkenhuysen M, Ye Z, Dudman JT, Dutta B, Hantman AW, Harris KD, Lee AK, Moser EI, O’Keefe J, Renart A, Svoboda K, Häusser M, Haesler S, Carandini M, Harris TD. Neuropixels 2.0: A miniaturized high-density probe for stable, long-term brain recordings. Science. 2021;372(6539). PMCID: PMC8244810

18. Juavinett AL, Bekheet G, Churchland AK. Chronically implanted Neuropixels probes enable high-yield recordings in freely moving mice. Elife. 2019;8:e47188. PMCID: PMC6707768

19. Bimbard C, Takács F, Catarino JA, Fabre JMJ, Gupta S, Lenzi SC, Melin MD, O’Neill N, Orsolic I, Robacha M, Street JS, Teixeira J, Townsend S, Beest EH van, Zhang AM, Churchland AK, Duan CA, Harris KD, Kullmann DM, Lignani G, Mainen ZF, Margrie TW, Rochefort NL, Wikenheiser AM, Carandini M, Coen P. An adaptable, reusable, and light implant for chronic Neuropixels probes. bioRxiv. 2024;2023.08.03.551752. PMCID: PMC10418246

20. Daal RJJ van, Aydin Ç, Michon F, Aarts AAA, Kraft M, Kloosterman F, Haesler S. Implantation of Neuropixels probes for chronic recording of neuronal activity in freely behaving mice and rats. Nat Protoc. 2021;16(7):3322–3347. PMID: 34108732

21. Horan M, Regester D, Mazuski C, Jahans-Price T, Bailey S, Thompson E, Slonina Z, Plattner V, Menichini E, Toksöz I, Pinto SR, Burrell M, Varsavsky I, Dalgleish HW, Bimbard C, Lebedeva A, Bauza M, Cacucci F, Wills T, Akrami A, Krupic J, Stephenson-Jones M, Barry C, Burgess N, O’Keefe J, Isogai Y. Repix: reliable, reusable, versatile chronic Neuropixels implants using minimal components. 2024;

22. Melin MD, Khanal A, Vasquez M, Ryan MB, Churchland AK, Couto J. Large scale, simultaneous, chronic neural recordings from multiple brain areas. bioRxiv. 2024;2023.12.22.572441. PMCID: PMC10769364

23. Juavinett A, Bekheet G, Churchland A. Implanting and Recycling Neuropixels Probes for Recordings in Freely Moving Mice. Bio-protocol. 2020;10(3):e3503. PMCID: PMC7842404

24. Song Z, Alpers A, Warner K, Iacobucci F, Hoskins E, Disterhoft JF, Voss JL, Widge AS. Chronic, Reusable, Multiday Neuropixels Recordings during Free-Moving Operant Behavior. eNeuro. 2024;11(1):ENEURO.0245-23.2023. PMCID: PMC10849027

25. Bimbard C, Takács F, Catarino JA, Fabre JM, Gupta S, Lenzi SC, Melin MD, O’Neill N, Orsolic I, Robacha M, Street JS, Teixeira J, Townsend S, Beest EH van, Zhang AM, Churchland AK, Duan CA, Harris KD, Kullmann DM, Lignani G, Mainen ZF, Margrie TW, Rochefort NL, Wikenheiser AM, Carandini M, Coen P. An adaptable, reusable, and light implant for chronic Neuropixels probes. 2025;

26. Weber JN, Peterson BK, Hoekstra HE. Discrete genetic modules are responsible for complex burrow evolution in Peromyscus mice. Nature. 2013;493(7432):402–405. PMID: 23325221

27. Ebensperger LA, Blumstein DT. Sociality in New World hystricognath rodents is linked to predators and burrow digging. Behav Ecol. 2006;17(3):410–418.

28. Makin DF, Agra E, Prasad M, Brown JS, Elkabets M, Menezes JFS, Sargunaraj F, Kotler BP. Using Free-Range Laboratory Mice to Explore Foraging, Lifestyle, and Diet Issues in Cancer. Frontiers Ecol Evol. 2021;9:741389.

29. Eilam D, Golani I. Home base behavior of rats (Rattus norvegicus) exploring a novel environment. Behav Brain Res. 1989;34(3):199–211. PMID: 2789700

30. Meng X, Chen P, Veltien A, Palavra T, Veld SI, Grandjean J, Homberg JR. Estimating foraging behavior in rodents using a modified paradigm measuring threat imminence dynamics. Neurobiol Stress. 2024;28:100585. PMCID: PMC10661863

31. Windolf C, Yu H, Paulk AC, Meszéna D, Muñoz W, Boussard J, Hardstone R, Caprara I, Jamali M, Kfir Y, Xu D, Chung JE, Sellers KK, Ye Z, Shaker J, Lebedeva A, Raghavan R, Trautmann E, Melin M, Couto J, Garcia S, Coughlin B, Elmaleh M, Christianson D, Greenlee JDW, Horváth C, Fiáth R, Ulbert I, Long MA, Movshon JA, Shadlen MN, Churchland MM, Churchland AK, Steinmetz NA, Chang EF, Schweitzer JS, Williams ZM, Cash SS, Paninski L, Varol E. DREDge: robust motion correction for high-density extracellular recordings across species. Nat Methods. 2025;22(4):788–800. PMID: 40050699

32. Buccino AP, Hurwitz CL, Garcia S, Magland J, Siegle JH, Hurwitz R, Hennig MH. SpikeInterface, a unified framework for spike sorting. eLife. 2020;9:e61834. PMCID: PMC7704107

33. Pachitariu M, Sridhar S, Stringer C. Solving the spike sorting problem with Kilosort. Biorxiv. 2023;2023.01.07.523036.

34. Xiang Y, Zhao Y, Cheng T, Sun S, Wang J, Pei R. Implantable Neural Microelectrodes: How to Reduce Immune Response. ACS Biomater Sci Eng. 2024;10(5):2762–2783. PMID: 38591141

35. Hermann JK, Capadona JR. Understanding the Role of Innate Immunity in the Response to Intracortical Microelectrodes. Crit Rev Biomed Eng. 2018;46(4):341–367. PMCID: PMC6391891

36. Zhao S, Tang X, Tian W, Partarrieu S, Liu R, Shen H, Lee J, Guo S, Lin Z, Liu J. Tracking neural activity from the same cells during the entire adult life of mice. Nat Neurosci. 2023;1–15. PMID: 36804648

37. Birman D, Yang KJ, West SJ, Karsh B, Browning Y, Siegle JH, Steinmetz NA. Pinpoint: trajectory planning for multi-probe electrophysiology and injections in an interactive web-based 3D environment. 2023;

38. Jones EAA. Assembly: Chronic recoverable Neuropixels in mice v2. 2023;

